# Detection of Macrobrachium rosenbergii golda virus (MrGV) in giant river prawn larvae across Asia through NCBI sequence read archive (SRA) data mining

**DOI:** 10.1101/2025.08.13.669918

**Authors:** Chantelle Hooper, Ronny van Aerle, David Ryder, Nicola M. Coyle

## Abstract

Macrobrachium rosenbergii golda virus (MrGV) was first characterised in *Macrobrachium rosenbergii* larvae in 2020, associated with mass mortalities in multiple Southern Bangladesh prawn hatcheries. MrGV has since been detected in two metatranscriptomic datasets from *M. rosenbergii* in China and in relation to a larval mortality event in India. The major objective of this study was to further characterise the geographical spread of the virus by mining the NCBI Sequence Read Archive (SRA) database. Utilising a new database, Logan, generated from assembling each SRA dataset, and a custom Snakemake pipeline, we discovered MrGV sequence data in *M. rosenbergii* SRA datasets from China, Thailand, and India, and determined that presence and relative abundance of MrGV is mostly associated with the larval life stage of *M. rosenbergii*. These results provide insights into the current prevalence of MrGV globally, suggest the life stages of prawn that should be screened to prevent spread of the virus, and demonstrate how the Logan database can be used to inform epidemiological studies.

## 2 Introduction

The giant river prawn, *Macrobrachium rosenbergii*, is a key aquaculture species cultivated in tropical and sub-tropical regions around the globe, but has suffered production issues due to multiple diseases (Hooper et al., 2022). Macrobrachium rosenbergii golda virus (MrGV) was first characterised in larval stages of *M. rosenbergii* in 2020, associated with mortalities in multiple Southern Bangladesh prawn hatcheries (Hooper et al., 2020). Two metatranscriptomic datasets of *M. rosenbergii* postlarvae and juveniles/adults from China have detected the presence of the virus, however disease or poor condition was only noted in one animal (Meng et al., 2022;Dong et al., 2024). During the course of this study, an investigation into large-scale mortality events in *M. rosenbergii* hatcheries in India during July - September 2024 found MrGV to be the causative agent (Paul et al., 2025). Despite these findings, the extent of MrGV presence in species of *Macrobrachium* globally is largely unknown.

MrGV falls within the *Nidovirales* order of enveloped positive-sense single stranded RNA viruses. Nidoviruses have a wide host range, including both vertebrates and invertebrates (ICTV, 2012). Invertebrate-infecting nidoviruses are a smaller group of viruses compared to those able to infect vertebrates (including SARS-CoV-2), however their diversity is becoming better known through metatranscriptomic viral discovery studies (Shi et al., 2016;Chang et al., 2021). Viruses within the *Roniviridae* family are some of the best recognised invertebrate-associated nidoviruses, comprising the yellow head viruses (YHV) (Chantanachookin et al., 1993), of which yellow head virus genotype 1 (YHV-1) is notifiable to the World Organisation for Animal Health (WOAH), due to its association with mass mortalities of penaeid shrimp (Walker et al., 2021;WOAH, 2024). Since its characterisation, MrGV has been formally recognised and named by the International Committee of Taxonomy of Viruses (ICTV) as *Nimanivirus lahi*, within the newly erected second genus of *Roniviridae, Nimanivirus* (Subgenus: *Marovirus*) (Gorbalenya et al., 2021).

The SRA database has expanded almost exponentially as more studies produce short read metagenomic, transcriptomic, and metatranscriptomic data to answer specific research questions. Previously, searching the SRA database for a target sequence to address various research questions required huge resources in time, storage and computing power, making screening of SRA datasets largely inaccessible to most users. By utilising massive cloud computing resources to carry out an SRA-wide genome assembly, the Logan dataset of DNA and RNA sequences (Chikhi et al., 2024) has reduced the redundancy and data volume of the SRA datasets, allowing these data to be searched at more efficiently and at an affordable cost to the user.

In this study we mined the *Macrobrachium* SRA database up to December 2023 for MrGV sequences, using Logan assembled SRAs to 1) further characterise the geographical spread of the virus, 2) determine if MrGV sequence type is associated with geographical location using phylogenetics, and 3) determine the species and life stages MrGV sequence is present in.

## 3 Methods

### 3.1 Selecting Sequence Read Archive (SRA) assessions and mapping Logan contigs

Logan assemblies of transcriptomes for all species of *Macrobrachium* deposited to the NCBI SRA database prior to 10/12/2023 were selected to be mined for MrGV sequences. The SRA accession numbers, their associated species, life stage and country of origin are outlined and summarised in Supplementary Tables 1 and 2. Logan-assembled contigs from release v1 were mapped to a MrGV reference genome (accession NC_076908) using minimap2 v2.28 with default parameters (Li, 2018) within a custom Snakemake (Mölder et al., 2021) v8.20.3 pipeline (logan-screen- accessions v1.0.0, DOI: 10.5281/zenodo.16418927). Samples were considered MrGV positive if they had at least 90% of bases covered by at least one contig, calculated using SAMtools v1.21 (Danecek et al., 2021).

### 3.2 Relative abundance of MrGV reads

To calculate the relative abundance of MrGV within SRA datasets containing Logan-assembled contigs that mapped to the reference genome, raw reads from the MrGV-positive SRA datasets were mapped to the MrGV reference genome using minimap2 v2.28, as above. To compare abundance between SRA datasets, the presence of MrGV was calculated as MrGV reads per million and transformed using the inverse hyperbolic sine (asinh) function. Relative abundance plots were generated with the ggplot2 R package (Wickham, 2016).

### 3.3 Construction of consensus MrGV genome sequences

Reads from biological replicates of SRA datasets with over 90% coverage of the reference genome were combined and mapped to the MrGV reference genome using minimap2, as described above, to increase coverage and enhance the likelihood of identifying small nucleotide polymorphisms (SNPs). The combined SRA datasets are detailed in Supplementary Table 3. A consensus sequence from the mapped reads was generated using SAMtools v1.21 (Danecek et al., 2021) in simple consensus mode, with call fraction set to 0.2. Combined raw reads from biological replicates (where they existed) were remapped to the resulting SAMtools-generated consensus sequence with minimap2, as above, and SNPs were called with Snippy v4.6.0 (Seemann, 2015), with default settings. A consensus genome with IUPAC codes at variable sites within each SRA dataset was generated from the raw output from snippy with the BCFtools v1.9 package (Danecek et al., 2021), setting the quality of mapping to greater than or equal to 100, minimum coverage to 30, and the minimum proportion for a variant to be called to 0.1. For MrGV genomes already deposited to NCBI, the associated SRA sequences (where present) were downloaded, SNPs were called, and consensus sequences with IUPAC codes were generated, as above.

### 3.4 Phylogenetic Analysis

A multiple sequence alignment (MSA) of complete MrGV genomes with degenerative nucleotide positions in IUPAC format and publicly available MrGV genomes was produced with MAFFT (Katoh and Standley, 2013) using the L-ins-I algorithm. A Bayesian consensus tree was constructed from the MSA using MrBayes v3.2.7 (Ronquist et al., 2012) on the CIPRES Science Gateway (Miller et al., 2010). The tree was constructed using two separate MC^3^ runs, carried out for two million generations using one cold and three hot chains. The first 500,000 generations were discarded as burn-in, and trees were sampled every 1000 generations. The resulting tree was midpoint rooted. A second Bayesian consensus tree, based on the translated sequence of ORF3, was generated as above.

## 4 Results

### 4.1 Macrobrachium rosenbergii golda virus (MrGV) presence in *Macrobrachium* spp. SRA datasets

A total of 965 *Macrobrachium* SRA datasets derived from RNA were screened for the presence of MrGV: 483 *M. rosenbergii*, 466 *M. nipponense*, six *M. australiense*, four *M. cancinus*, two *M. novaehollandiae*, two *M. tolmerum*, one *M. olfersii*, and one *M. koombooloomba*. MrGV was only found to be present in *M. rosenbergii* SRA datasets, appearing across all life stages, but with a notable preference for the larval life stage. Reads mapped to MrGV in 27% of *M. rosenbergii* embryo SRA datasets (n = 88), 61% of larval SRA datasets (n = 150), 41% of postlarvae SRA datasets (n = 41), and 7% of juvenile/adult SRA datasets (n = 201) (Figure 1A). MrGV also appeared to be present at a higher relative abundance in larvae compared to other life stages (Figure 1B). Larvae had a median relative abundance of 6,013 MrGV reads/million, while embryos, postlarvae and adults had median relative abundances of 3.73, 510.92 and 0.021 MrGV reads/million, respectively. Coverage across the MrGV genome was not uniform, with the majority of reads mapping to the 3’ end of the virus (Supplementary Figure 1).

**Figure 1:**
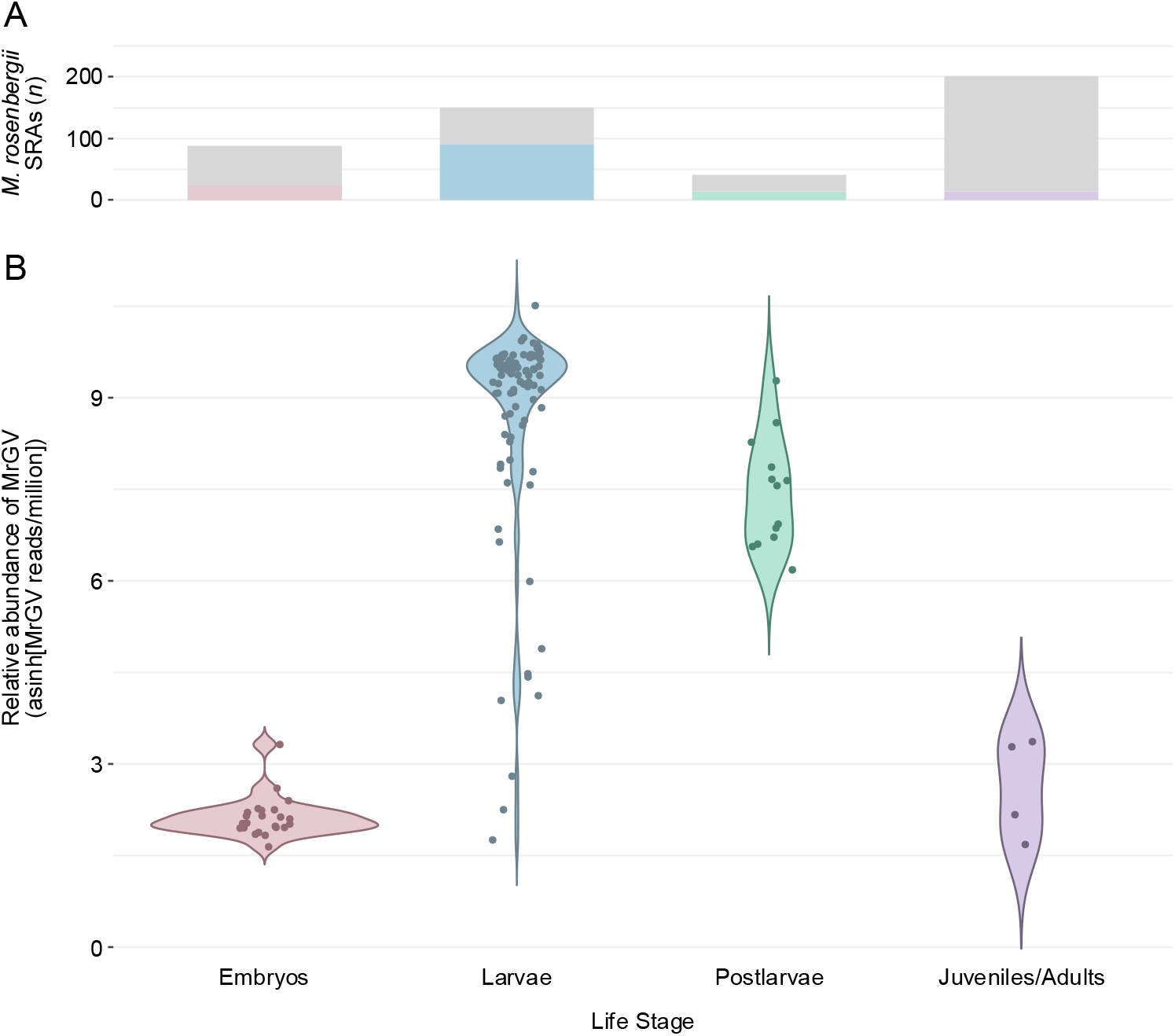
Presence of Macrobrachium rosenbergii golda virus (MrGV) in *Macrobrachium rosenbergii* sequence read archive (SRA) datasets by life stage. (A) Bar plot showing the number of *M. rosenbergii* SRA datasets containing MrGV sequences by life stage. Grey bars represent the absence of MrGV in Logan-assembled SRA contigs, and coloured bars represent the presence of MrGV. (B) Violin plot showing the relative abundance of MrGV reads (reads/million, transformed with the inverse hyperbolic sine function) within *M. rosenbergii* SRA datasets published prior to 10^th^ December 2023. SRA datasets with fewer than one MrGV read per million were excluded from the plot.

### 4.2 The relative abundance of MrGV increases over larval development

SRA datasets that appear to relate to a timed developmental study on larval stages of *M. rosenbergii* at different salinities (BioProjects PRJNA864119 and PRJNA891247) showed an increase in MrGV read presence over the first 20 days of development. These BioProjects, which lack associated publications, sampled larvae in triplicate at 0-, 6-, 12-, 24-, and 48-hours post-hatch, followed by sampling on days 5, 10, and 20. Larvae were developed in salinities of 5, 15, and 25, with the units presumed to be parts per thousand (‰), although no units were stated in the SRA metadata. As there is no associated publication for these data, clinical signs of disease and mortality data were not available.

When transformed with the inverse hyperbolic sine function and plotted (Figure 2), MrGV relative abundance follows the same pattern for larvae developed at each salinity, peaking at 10-15 days post-hatch, before plateauing. Average MrGV relative abundance stayed below 10 MrGV reads/million until 2-days post-hatch, where average MrGV reads increased to between 200-625 reads/million. Generally, the relative abundance MrGV reads continued to increase to approximately 7,000 reads/million between days 5-10, which was maintained until the end of the experiment at 20-days post-hatch.

**Figure 2:**
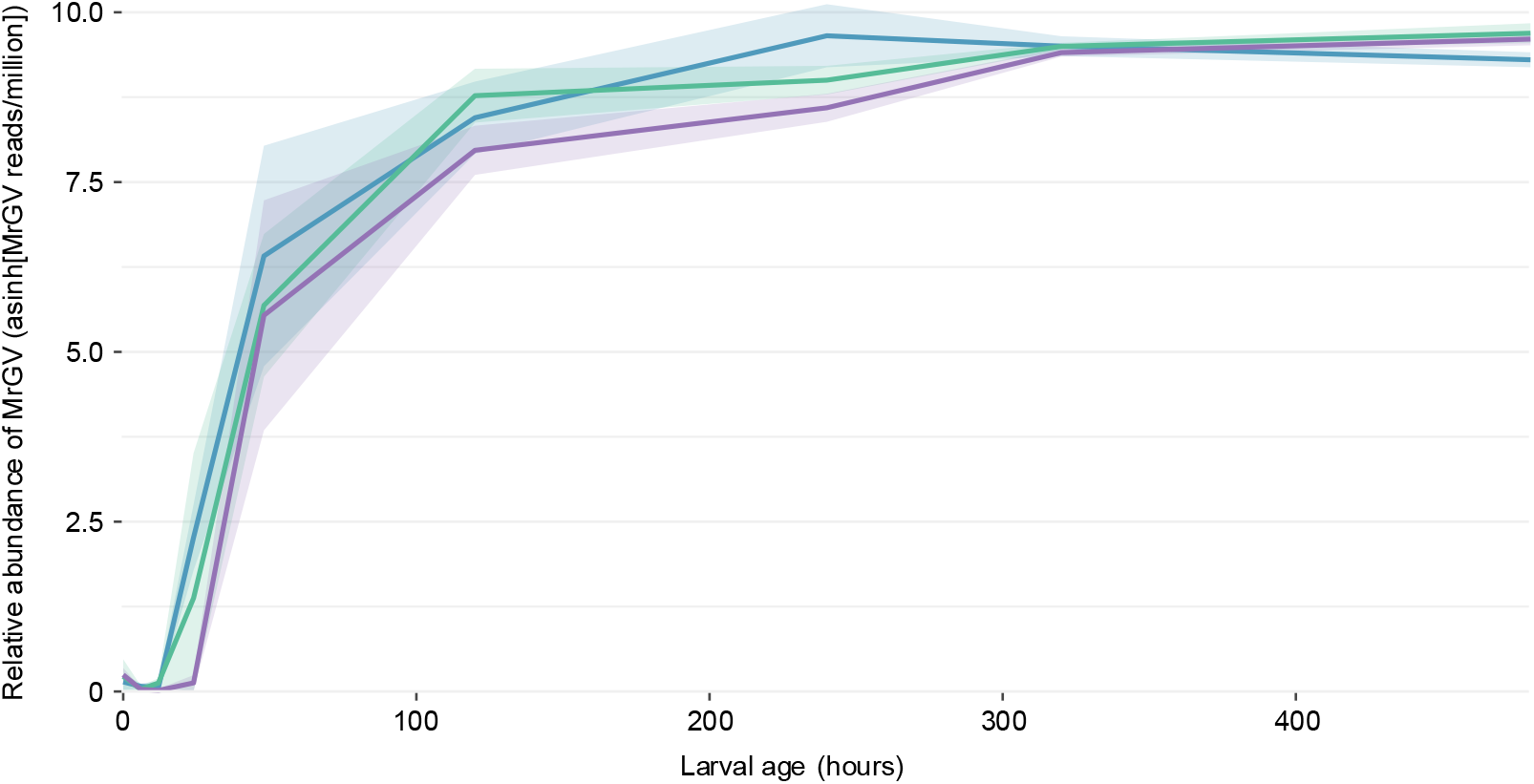
Line plot showing the change in Macrobrachium rosenbergii golda virus (MrGV) relative abundance over 20 days of larval development at different salinities (NCBI BioProjects PRJNA864119 and PRJNA891247). Salinity units from the SRA metadata were presumed to be in parts per thousand (‰). Each coloured line represents development at a different salinity, with the 95% confidence interval depicted by a ribbon in a lower transparency of the same colour: Blue represents development at 5‰, green represents development at 15‰, and purple represents development at 25‰.

MrGV was also present in *M. rosenbergii* embryos. Embryo SRA datasets with reads that mapped to MrGV originated from a single BioProject (PRJNA890910). This BioProject, submitted to NCBI by the same institute and at the same time as the larval salinity challenge, appears to study the transcriptome of *M. rosenbergii* over the course of embryonic development, however no associated publication could be found and we were unable to link the embryos to the salinity-challenged larvae. MrGV reads were present in all replicates (three per developmental stage) for all stages of embryonic development: first and second cleavage, blastula, gastrula, egg nauplius, egg metanauplius, egg protozoea and egg zoea stages. However, the number of MrGV reads did not change over time and remained at a consistently low relative abundance, averaging between 2 and 8 MrGV reads/million over the course of the study.

### 4.3 MrGV sequence type varies by location

A total of 88 SRA datasets were used to assemble 30 novel MrGV genomes. Reads from biological replicates within the same BioProject were merged in order to increase MrGV coverage and the ability to call SNPs. A further six MrGV genomes were obtained from NCBI GenBank. Of the 36 MrGV genomes, 31 originated from China, two from Thailand, two from India, and one from Bangladesh. The 31 genomes from China originated from three provinces: two neighbouring provinces in the Southeast of China, Guangdong and Guangxi, and one province in the East of China, Jiangsu. None of the Logan contigs originating from *M. rosenbergii* SRA datasets from Israel, Malaysia or Vietnam mapped to the reference MrGV genome.

A midpoint rooted Bayesian tree constructed from all complete MrGV genomes demonstrated that MrGV from each location branched separately from one another (**Figure 3**). MrGV from Thailand, India and Bangladesh branched as distinct, full supported clades. MrGV from each province in China also branched separately, with full support for each clade. MrGV from the two geographically close provinces, Guangdong and Guangxi, branched as sister clades, while MrGV from the geographically distant province, Jiangsu, branched separately. Jiangsu is the only location where SRA datasets exist covering multiple years; despite this temporal difference, MrGV genomes still branch together. MrGV genome MW590703 had no associated SRA dataset, therefore the distance between this genome and other genomes from Jiangsu could be partially explained by the lack of degenerate bases within the genome, as variants could not be called.

**Figure 3:**
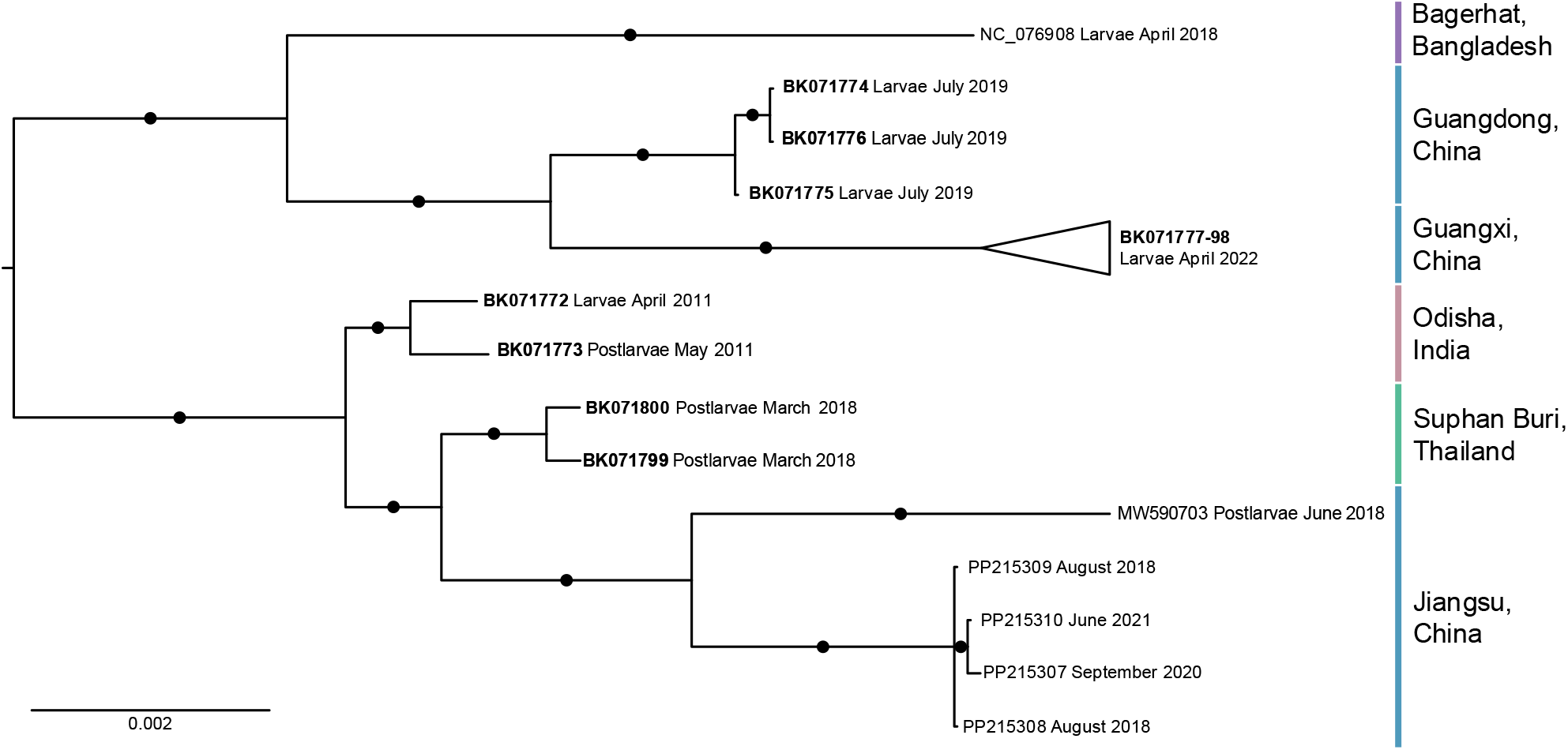
Midpoint rooted Bayesian consensus tree constructed from 36 full genome sequences from Macrobrachium rosenbergii golda virus. Circles on branches represent posterior probabilities of 1, and bold accession numbers represent genomes assembled in this study.

The nucleotide differences that cause MrGV from different locations to branch separately occurred across the MrGV genome, with the majority of these nucleotide differences (86.27%) not causing changes in the resulting amino acid sequence. Base changes that caused alterations in amino acid sequence accounted for 13.68, 11.82, 10.00, and 21.18% of mutations in ORFs 1a, 1b, 2 and 3, respectively. Amino acid changes in the key nidovirus protein motifs within ORF1b were low, with no mutations that conferred an amino acid change in the S-adenosylmethionine (SAM)-dependent N7-methyltransferases (N-MT) motif, and only one in the nidovirus RdRp-associated nucleotidyltransferase (NiRAN) and zinc-binding domain (ZBD) motifs (Supplementary Figure 2). A large non-cytoplasmic domain at the 5’ end of ORF3, thought to encode one of the envelope glycoproteins, contained 14 sites in the 5’ region of the domain that resulted a change in amino acid sequence. Given that the metadata associated with the SRA datasets were limited, we were unable to determine if these differences in glycoprotein sequence conferred a difference in the virulence or pathogenicity of MrGV. Variation in the 5’ untranslated region (UTR) of the MrGV genomes was not assessed due to low sequence coverage, however the 3’ UTR, known to contain the presence of a secondary RNA structure nidoviruses (Dreher, 1999), was well conserved between MrGV genomes from all locations, with only two nucleotide differences present (compared to the reference sequence) in MrGV from Guangxi, China.

A second Bayesian consensus tree was constructed from a multiple sequence alignment of the translated sequence of ORF3 for all SRAs (or concatenated biological replicates) where a full ORF3 sequence was present. This tree (**Figure 4**) had more paraphyletic clades when compared to the tree constructed from full genomes (**Figure 3**), and in general, had lower posterior probabilities. However, as in the tree generated from full genomes, samples from Guangdong and Guangxi branched separately from other MrGV samples with maximal support, and within this clade they branched as highly supported sister clades. Samples from Jiangsu also formed a separate clade, but with lower support (0.82 posterior probability) than in the tree constructed from full genomes.

**Figure 4:**
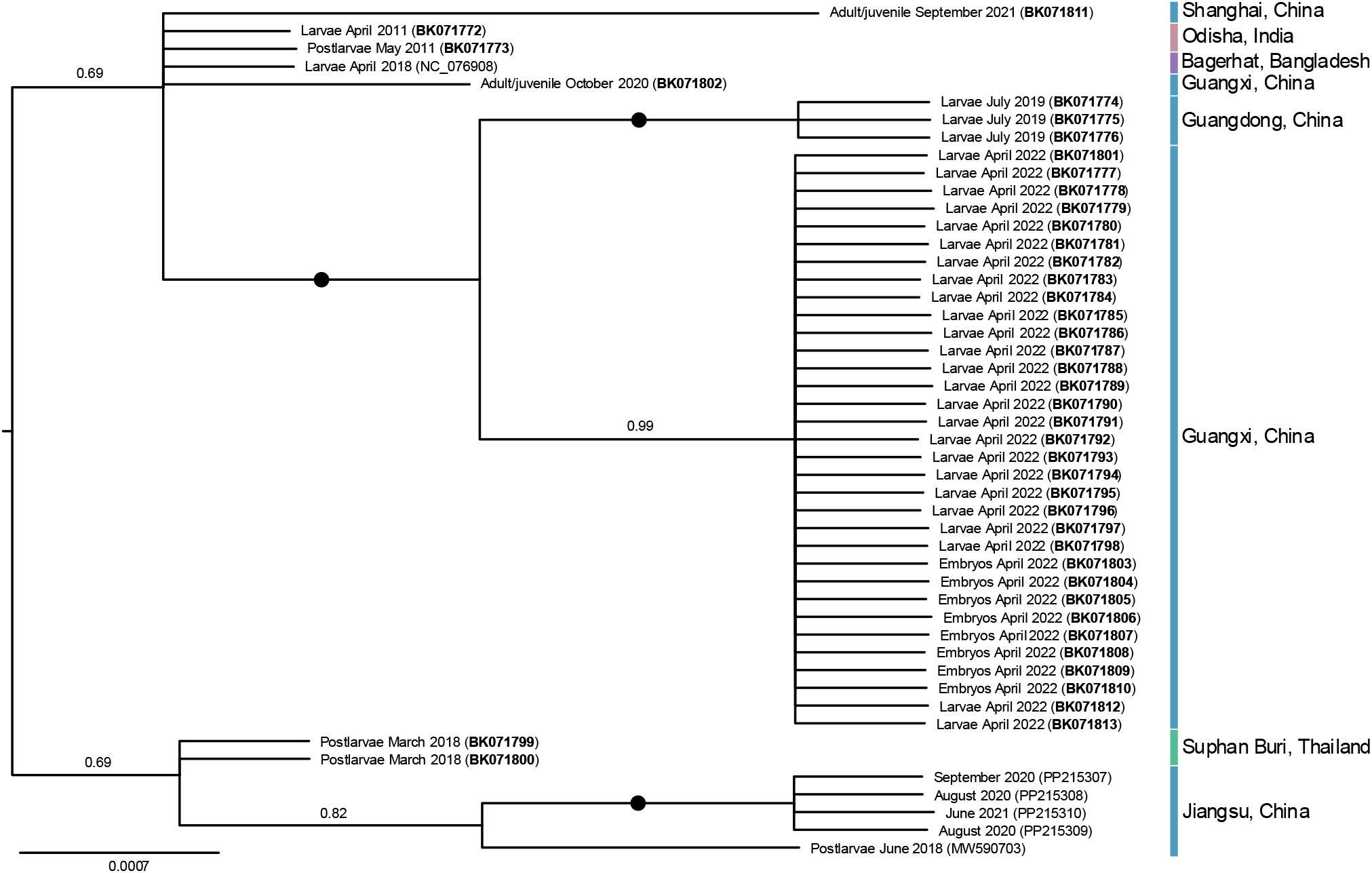
Midpoint rooted Bayesian consensus tree constructed from translated ORF3 sequences (n = 48) from Macrobrachium rosenbergii golda virus. Circles on branches represent posterior probabilities of 1, and bold accession numbers represent sequences generated in this study.

## 5 Discussion

In this study we found Macrobrachium rosenbergii golda virus (MrGV) present in *M. rosenbergii* SRA datasets from multiple locations in Asia. Previously, MrGV had only been reported from Bangladesh (Hooper et al., 2020), one province in China (Jiangsu) (Meng et al., 2022;Dong et al., 2024), and India (Paul et al., 2025). However, we found MrGV sequence data in prawns from Thailand and in an additional two provinces in the southeast of China; we also find MrGV in SRAs from India from 2011. In addition to the geographical distribution of MrGV, we determined that presence and high relative abundance of MrGV is highly associated with the larval life stage of *M. rosenbergii* and is apparently absent from other species within the *Macrobrachium* genus, although SRA datasets for larval stages of other *Macrobrachium* species were limited (Supplementary Table 2). Despite adult and juvenile SRAs making up the largest proportion of *M. rosenbergii* datasets, very few animals had Logan contigs that mapped to the MrGV genome and MrGV relative abundance was low.

Mass mortalities of *M. rosenbergii* larvae in Bangladesh hatcheries first occurred in 2011 (Hooper et al., 2020). Although no samples of moribund larvae exist to confirm MrGV infection in these early mortalities, the clinical signs of disease were identical to MrGV infection. Here, we show that MrGV was also present in larvae and postlarvae (SRA accessions DRR023219 and DRR023253, respectively) from Odisha, India, at the same time that the mortalities started to occur in Bangladesh, however the metadata associated with the larvae and postlarvae from India state that these animals were healthy. Interestingly, the study that describes MrGV as the causative agent of mass mortalities in *M. rosenbergii* hatcheries in India in 2024 (Paul et al., 2025) was from the same institute in India that submitted the SRA datasets in 2011 that we found to contain MrGV. Future investigation should be directed towards understanding how MrGV was introduced into hatcheries in both Bangladesh and India in 2011, and determine why, if MrGV has been present in Odisha since 2011, have mortalities only started occurring recently. Particular attention should be given to understanding whether *M. rosenbergii* genetics or rearing conditions play a role in the susceptibility of larvae to MrGV.

The study that initially characterised MrGV suggested that additional work was required to determine how the virus was entering hatchery systems (Hooper et al., 2020). In this study, we find MrGV reads in SRA datasets for all stages of *M. rosenbergii* embryo development, suggesting that vertical transfer of MrGV is a possibility. Paul et al. (2025) also found MrGV to be present at low abundance in *M. rosenbergii* eggs collected from adults, therefore collection of wild and/or unscreened berried female *M. rosenbergii* as broodstock for hatcheries may be a route of transfer of MrGV into hatcheries. Stress has long been portrayed as a major facilitator in the progression of disease in aquaculture (Meyer, 1991), and the stress of culture conditions may be a factor in the susceptibility of larvae to MrGV infection. *M. rosenbergii* larvae perform best at a salinity of 13‰ (Wei et al., 2021), hence the salinity stress delivered by culturing *M. rosenbergii* larvae at salinities higher or lower to this value, e.g. those used in BioProjects PRJNA864119 and PRJNA891247 (**Figure 2**), may predispose animals to infection with MrGV.

The changes in relative abundance of MrGV over larval development provides knowledge of the point in larval development when MrGV becomes abundant within prawn larvae. Due to the pooling of larvae from multiple timepoints in Hooper et al. (2020), it was only possible to determine that MrGV was lower in abundance 1-4 days post-challenge than MrGV to 5-10 days post-challenge. A similar trend was observed in the salinity challenged larvae (**Figure 2**), with the initial increase in MrGV relative abundance associated with larvae 2-days post-hatch, and MrGV relative abundance peaking between days 5-10 post-hatch before plateauing. The increase in MrGV relative abundance coincided with the time in larval development during which *M. rosenbergii* larvae are typically within the zoea III stage (Ling, 1969). Paul et al. (2025) reports that larval mortalities occur in hatchery settings from zoea III to zoea IX. Although the timed larval study (Figure 2) only demonstrates the relative abundance of MrGV up to 20 days post-hatch, so misses the later zoea stages, MrGV abundance was highest when larvae were likely in the zoea III-VII stages.

This study was limited by substantial data gaps and inconsistencies within the metadata associated with the SRA database. One of the most significant challenges was the ability to link publications to data from BioProject, BioSample, or SRA identifiers to fill these metadata gaps. Publication and dataset identifiers are not automatically connected by NCBI, and identifiers are often inconsistently cited or omitted from publications. Given these missing data, particularly those associated with the health status or clinical signs of disease of the animal or tissue that was sequenced, we were unable to associate the presence of *M. rosenbergii* with poor health or moribundity. The current study was also limited by the number of larval datasets available in the SRA database. As the Logan database only included data that were available in the SRA prior to the end of 2024, no assembled SRA sequences were available for larval stages of *Macrobrachium* species except for *M. rosenbergii*, and SRA datasets for postlarvae were limited to 45 datasets from *M. nipponense* and one from *M. australiense*. Given these limitations, we cannot confidently determine that the host range of MrGV is limited to *M. rosenbergii*, assuming that if it is able to infect other species of *Macrobrachium*, it would be primarily associated with the larval life stage.

A lack of SRA metadata regarding the health status of the host meant that it was not possible to associate any amino acid changes in key MrGV motifs to the infectivity of MrGV. Mutations in glycoprotein motifs in other nidoviruses have been shown to affect functional properties, including altering viral infectivity. Due to the large number of genomes and extensive patient data associated with infection with SARS-CoV-2, single mutations that conferred animo acid changes within the spike glycoprotein could be associated with increased or decreased binding to host receptors and subsequently linked to a change in infectivity (reviewed in Harvey et al. (2021)). To a lesser extent, this type of study has been carried out within the YHV complex of viruses, where only genotypes 1 and 2 have been associated with yellow head disease, while other genotypes persist at low levels in healthy animals (Wijegoonawardane et al., 2008). It has been suggested that variation in a critical cleavage site within the yellow head virus glycoprotein gp116 regulates the availability of viral glycoproteins, ultimately influencing whether the virus is able to cause chronic persistent infection or acute infection and disease (Wijegoonawardane et al., 2008). Conversely, even with a large deletion within the gp116 cleavage site in yellow head virus genotype 1b, the virus remains highly virulent and infectious, despite significant structural deformation that reduces its incorporation into virions (Sittidilokratna et al., 2009). Given this, information on the health status of animals associated with viral detection is essential to link predictions of virulence inferred from sequence changes to the ability of a pathogen to cause disease.

## 6 Summary

This study demonstrates how the Logan database can be utilised to rapidly and efficiently find target sequences within the SRA database, expanding the use of these publicly available sequencing datasets outside of their original intended purposes, for example using transcriptomic studies to find associated pathogens. Here, we used the Logan database to search for an emerging virus (Macrobrachium rosenbergii golda virus) in prawns and demonstrated how this can add valuable knowledge about the virus, including geographical spread, host range, and relative abundance, without the need for additional sampling or experimental infection. However, metadata associated with SRA datasets are typically inconsistently entered and poorly maintained. Without consistent, comprehensive metadata, the reliable interpretation of sequencing data remains challenging. Careful application of this approach, alongside improvements in metadata quality and accessibility, could uncover critical insights into pathogen biology, transmission and control. This type of data mining could add otherwise unknown data to epidemiological studies of emerging, reemerging, and rare pathogens globally, allowing the determination of spread of agents within and between populations, and govern which compartments of a system are most appropriate to screen to prevent pathogen spread.

## 8 Data availability

MrGV genomes generated in this study have been deposited to NCBI under accession numbers BK071772 - BK071801. Partial MrGV sequences (covering ORF3 only) are deposited under accession numbers BK071802 - BK071813. Details of the SRA datasets used to construct MrGV sequences are given in Supplementary Table 3.

