## Supplementary Figure for "Detection of Macrobrachium rosenbergii golda virus (MrGV) in giant river prawn larvae across Asia through NCBI sequence read archive (SRA) data mining"

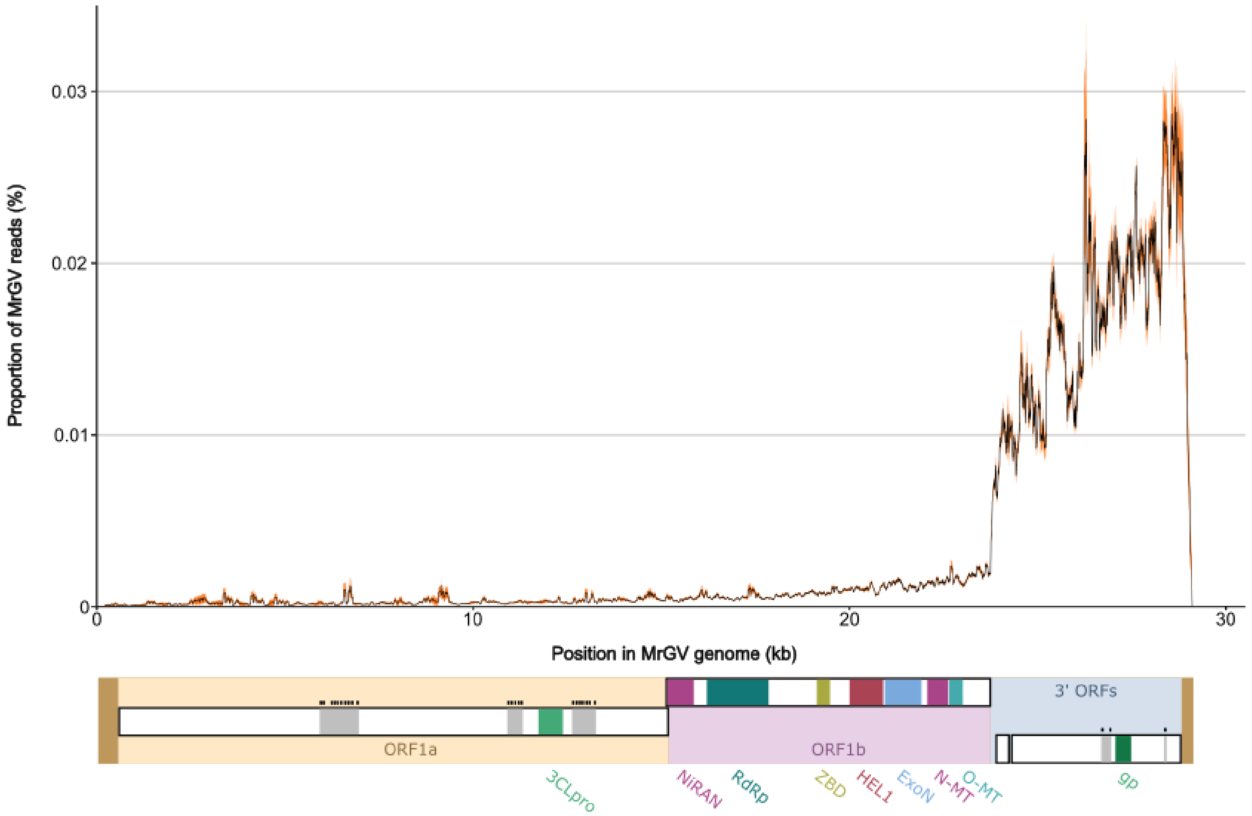


**Supplementary figure 1:** Graph showing the location of mapped reads across the MrGV genome from all SRAs ($n$ = 145). Orange ribbon represents the 90% confidence interval.

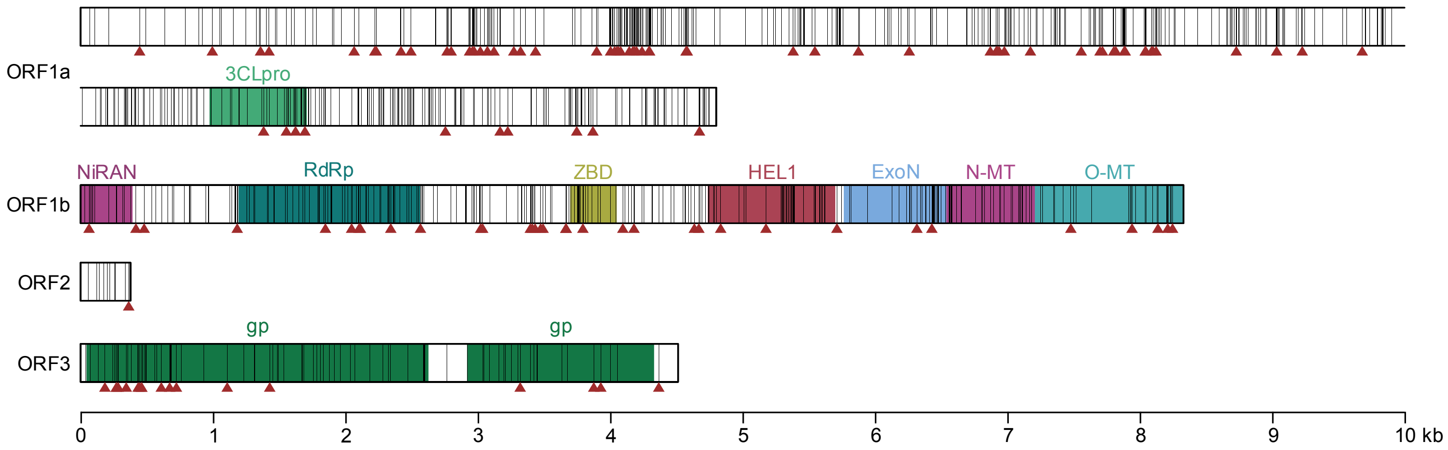


**Supplementary figure 2:** Location of sites within the coding regions of the MrGV genome that had a nucleotide change in at least one full MrGV genome. Vertical black lines depict all sites where a nucleotide difference was observed, and red arrows show where these nucleotide changes caused a change in amino acid sequence. Predicted protein motifs are a 3C-like protease (3CLpro), nidovirus RdRp-associated nucleotidyltransferase (NiRAN), RNA-dependent RNA polymerase (RdRp), zinc-binding domain (ZBD), superfamily 1 helicase (HEL1), 3'-5' exoribonuclease (ExoN), S-adenosylmethionine (SAM)-dependent N7- and 2'-O-methyltransferases (N-MT and O-MT, respectively) and glycoproteins (gp).
